# Artificially selecting microbial communities using propagule strategies

**DOI:** 10.1101/2020.05.01.066282

**Authors:** Chang-Yu Chang, Melisa L. Osborne, Djordje Bajic, Alvaro Sanchez

## Abstract

Artificial selection is a promising approach to manipulate the function of microbial communities. Here, we report the outcome of two artificial selection experiments at the microbial community level. Both experiments used “propagule” strategies, in which a set of the best-performing communities are used as the inocula to form a new generation of communities. In both cases, the selected communities are compared to a control treatment where communities are randomly selected. The first experiment used a defined set of strains as the starting inoculum, and the function under selection was the amylolytic activity of the consortia. The second experiment used a diverse set of natural communities as the inoculum, and the function under selection was the cross-feeding potential of the resulting communities towards a reference bacterial strain. In both experiments, the selected communities reached a higher mean and a higher maximum function than the control. In the first experiment this is caused by a decline in function of the control, rather than an improvement of the selected line. In the second experiment, the strong response of the mean is caused by the large initial variance in function across communities, and is the immediate consequence of the spread of the top-performing community in the starting group, whose function does not increase. Our results are in agreement with basic expectations of artificial selection theory, pointing out some of the limitations of community-level selection experiments which can inform the design of future studies.

## Introduction

Microbes form complex ecological communities consisting of large numbers of interacting taxa. The collective activity of all species within these communities can have important ecological and biotechnological impacts, for instance by affecting the health and life-history traits of their hosts (Fraune and Bosch 2010; Human Microbiome Project Consortium 2012; Santhanam et al. 2015; Hu et al. 2016; Berendsen et al. 2018), or by producing valuable products from waste materials (Bernstein and Carlson 2012; Shong et al. 2012; Minty et al. 2013; Wang et al. 2016). Manipulating community composition towards desirable collective functions, and away from undesirable ones, has become an important aspiration in microbiome biology and biotechnology (Mueller and Sachs 2015; Eng and Borenstein 2016; Rillig et al. 2016; Sheth et al. 2016). One approach has been to engineer communities from the bottom-up, by mixing species with known functionality towards a desired community function (Gilbert et al. 2003; Regot et al. 2011; Minty et al. 2013; Smith et al. 2013; Wang et al. 2016; Eng and Borenstein 2019). An important challenge of this approach is the combinatorial explosion in the number of interactions in the community, with high-order interactions posing significant challenges for the predictability of even mid-size consortia (Gould et al. 2018; Mickalide and Kuehn 2019; Sanchez-Gorostiaga et al. 2019; Senay et al. 2019). In addition, ecological and evolutionary processes can also act against the engineered community function (Goldman and Brown 2009).

The difficulty of engineering community functions from the bottom-up (Escalante et al. 2015) has motivated a surge of interest in evolutionary design (Bentley 1999) approaches, which treat the community as a unit of selection and explore the ecological landscape in search of consortia with desirable traits (Arias-Sánchez et al. 2019). This approach has been found to work *in silico* (Penn 2003,Williams and Lenton 2007a,b; Doulcier et al. 2019; Xie et al. 2019) and it has been attempted several times in the laboratory to optimize functions such as toxin removal (Swenson et al. 2000a), the manipulation of environmental pH (Swenson et al. 2000b), or the modulation of various host traits (Swenson et al. 2000b; Mueller and Sachs 2015; Panke-Buisse et al. 2015, 2016; Gopal and Gupta 2016; Mueller et al. 2016; Jochum et al. 2019). The results of these experimental studies have been mixed (Blouin et al. 2015; Arora et al. 2019), and it is becoming increasingly clear that the details of how exactly the “offspring” communities are generated from the “parental” communities can be critical for the success of this approach (Mueller et al. 2016; Raynaud et al. 2019).

One of these methods of community generation can be thought of as being akin (in a metaphorical sense) to “asexual” reproduction (Swenson et al. 2000b) and has been termed the “propagule method” (Swenson et al. 2000b). In this method, communities are ranked on the basis of the function of interest, and the top-ranked communities are selected to inoculate a new generation by taking a small aliquot and introducing it into fresh new habitats (Fig. 1A). Thus, each selected community acts as the single parental community that seeds a new crop of *N*>1 new “offspring” communities. This method has been used in at least three different studies to our knowledge (Swenson et al. 2000b; Arora et al. 2019; Raynaud et al. 2019). An alternative method has also been used and termed the “migrant pool method” (Swenson et al. 2000b). It relies on mixing together the highest performing communities, using the mixed pool as the inoculum from which the offspring generation will be seeded (Swenson et al. 2000b; Blouin et al. 2015; Panke-Buisse et al. 2015; Raynaud et al. 2019).

**Fig. 1.**
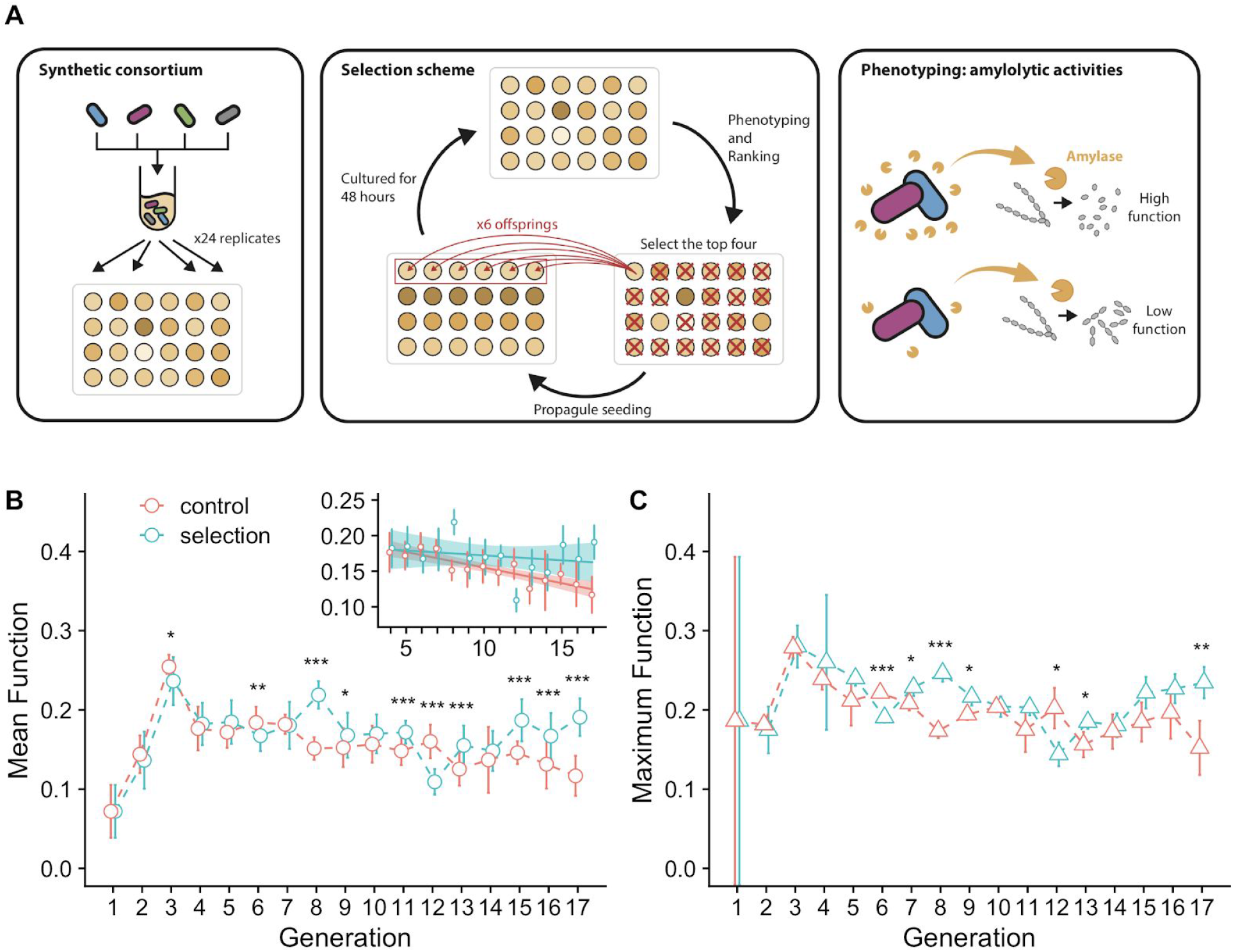
Artificial selection of an amylolytic consortium. **A.** Experimental scheme. **B**. Community function as the fraction of starch hydrolyzed by the community filtrate. Green circles represent the mean functions (± 1 SD) of the 24 communities in the selection line. Red open circles represent the random control line. Asterisks indicate generations in which the community functions differ significantly between selection and control lines (Welch’s two-sample t-test. *** indicates p < 0.001, ** indicates p<0.01, * indicates p<0.05). Inset shows the linear regression of mean community function to the rounds of selection (generations), starting right after the beginning of the stable phase at generation 4 (slope = −0.0013, P = 0.44 for selection line; slope = −0.004, P < 0.001 for control line). **C**. Highest community function in each generation for both the selection (green) and the control lines. Green triangles represent the mean of the triplicate measurement of the best performed community (± 1 SE, N=3) in the selection line. Red triangles represent the control line. Asterisks denote the same significant levels as in **B**.

The objective of this paper is to report the outcome of two independent experiments that used propagule strategies for community reproduction. We note that our intention is not to systematically and directly compare the outcomes of both experiments, as they differ in the amount of community variability in the starting community pool, in the experimental design, and in the function under selection. Despite these differences, some general patterns are found, and our goal is to make the results available to the community so they can be of use for the future design of new experiments and protocols. In the Results section we will limit ourselves to describe the two experiments and their outcomes, which we will then unpack in the Discussion.

## Results

### Artificial selection of an amylolytic consortium

Designing communities for efficient biodegradation has long been an aspiration of synthetic ecology and community engineering (Gilbert et al. 2003; Yoshida et al. 2009; Zanaroli et al. 2010; Piccardi et al. 2019), and it has also been a problem of interest in artificial community-level selection and microbial bioprospecting (Swenson et al. 2000a; Zanaroli et al. 2010; Cortes-Tolalpa et al. 2016; Eng and Borenstein 2019). In particular, the hydrolysis of complex polymers into smaller molecular weight molecules (which can be then metabolized into useful byproducts by other community members) is often a limiting step in bioconversion (Minty et al. 2013; Kim et al. 2014; Yang et al. 2016). As a proof of concept, we set out to apply artificial selection to find a community with an enhanced ability to hydrolyze starch. While the hydrolysis of starch is not in general as challenging as that of other polymers (e.g. cellulose), it is also of value for industrial applications (de Souza and de Oliveira Magalhães 2010; Xu et al. 2016). In our experiments, we used soluble starch as a substrate, as its hydrolysis has been recently characterized as an ecological function at the community-level (Sanchez-Gorostiaga et al. 2019) that is relatively straightforward and inexpensive to characterize in high-throughput (Fuwa 1954).

Our starting point was a set of strains of soil bacteria whose amylolytic ability we have recently characterized (Sanchez-Gorostiaga et al. 2019), including *B. subtilis, B. licheniformis, B. megaterium*, and *P. polymyxa*. All communities in the metacommunity were inoculated from the same pool of species, which consisted of the 4 strains listed above mixed in equal ratios (as measured by OD620nm). To begin the experiment, all wells of a 24-well plate were inoculated from the species pool and allowed to grow for 48 hours as described in Methods. After that time, a 500uL volume of each culture was filtered-sterilized and used to assay for enzymatic starch hydrolysis (Methods). The function of each community represents the fraction of starch degraded over a 10min incubation period by 20uL of its filtered medium at 30°C (Methods). The four communities with the highest function were set aside, and each of them was used as the inoculum for a new generation of communities (6 communities were seeded from each; Fig. 1A, Methods). As a control, a random treatment was started where 4 communities were randomly selected in each generation as the inocula to seed the next generation (Fig. 1A, Methods).

The mean amylolytic activity for both treatments is shown in Fig. 1B. Both treatments are initially indistinguishable and follow identical trends, with an initial rise in the mean function that ends on the third generation, followed by a stable phase where the function varied little over time. The selected treatment had a higher mean function than the control in the last three days of the experiment (Welch’s two-sample t-test, p<0.01) (Fig. 1B). However, this is not caused by an increase in the mean function in response to selection, which in fact remains stable after the stabilization phase starts on generation 4 (Mann-Kendall Test *τ* = - 0.14, P=0.47), but rather by a statistically significant decline in the mean function of the control line over time (Mann-Kendall Test *τ* = −0.67, p<0.001)(Fig. 1B; Inset). The same pattern was observed when we measured the function of the top community for each treatment: The maximum function in the community is also stable in the artificial selection line (Mann-Kendall Test *τ* = −0.25, P=0.21) after the third generation, but it exhibits a statistically significant decline over time in the control (Mann-Kendall Test *τ* = −0.58, P=0.004) (Fig. 1C). The lack of response to artificial selection in our experiments can be explained by the absence of a significant heritable component of the community-level functional variance (Goodnight 2000) (h^2^=0.05, P=0.3, estimated by regressing the offspring to the parent functions following (Blouin et al. 2015)).

### Artificial selection of a cross-feeder community

A promising application of microbial community engineering (Wei et al. 2015; Jousset et al. 2016) is biocontrol. Biocontrol encompasses both the suppression of pathogen growth (Mendes et al. 2011; Berendsen et al. 2018) and the promotion of other (potentially beneficial) organisms or hosts. In the second experiment, we set out to use artificial community-level selection to find a community that would strongly promote the growth of a target organism (Fig. 2A). Our target organism was an *E. coli* strain that cannot grow aerobically on citrate (the only externally supplied carbon source in the growth medium) (Fig. 2A). By adding the byproducts released by a community to a monoculture of *E. coli*, we would be supplementing the medium with additional carbon sources, making it possible for *E. coli* to grow. The cross-feeding ability of a community can be quantified by measuring the yield of *E. coli* in the byproducts of that community. For internal consistency, this yield was normalized to that of an *E. coli* culture growing on glucose as the only carbon source on the same plate, incubator, media, etc. as the rest of experiment (Fig. 2A; Methods). This ratio represents the function under selection.

**Fig. 2.**
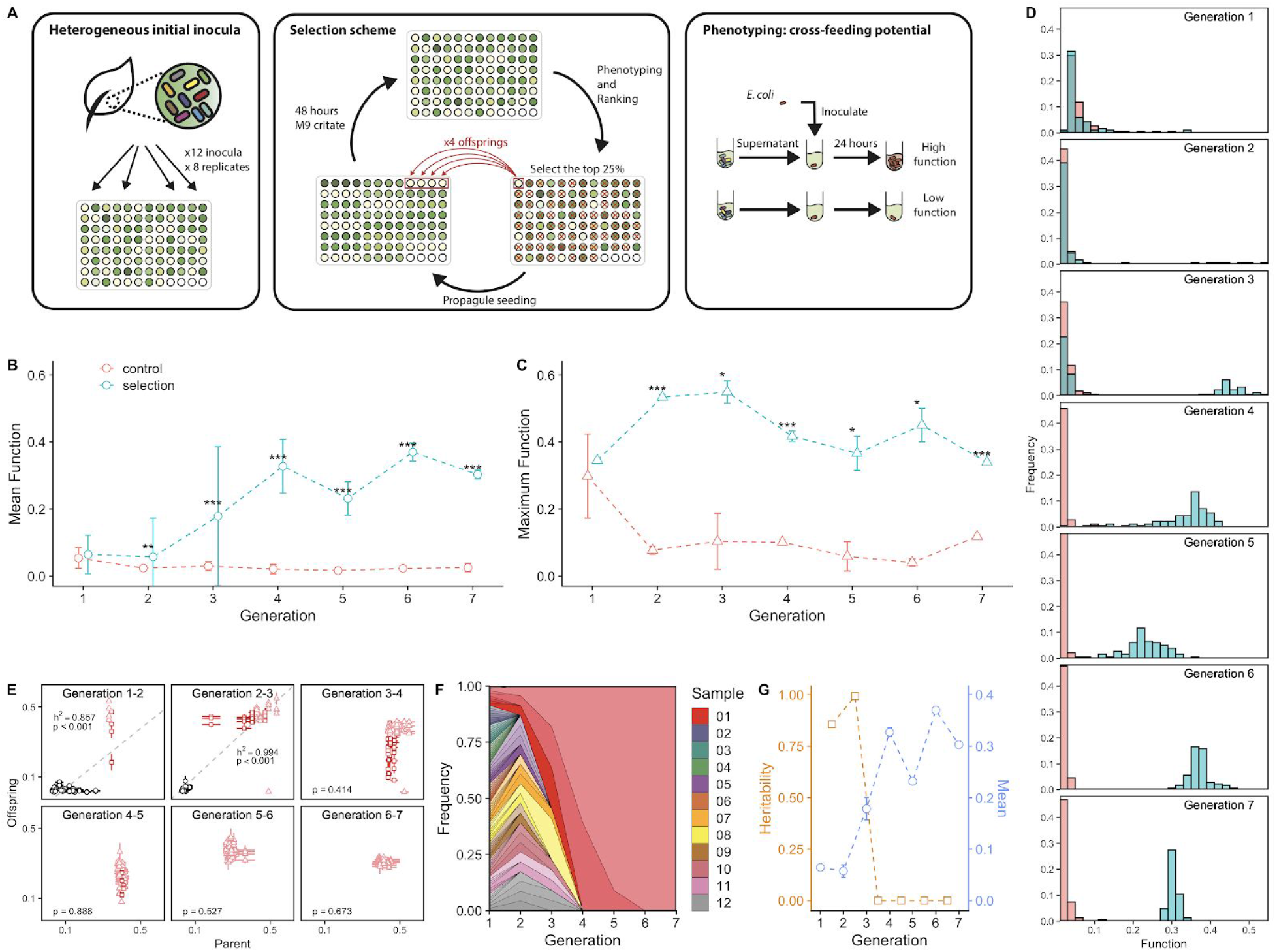
Artificial selection of cross-feeder communities. **A.** Experimental scheme: the artificial selection on the cross-feeding potentials of communities on a target organism *E. coli*. Cross-feeder communities were obtained from 12 leaf and soil samples and grown on citrate-M9 medium. Community function as the ratio of Optical Density at 620nm (OD_620_) of *E. coli* grown on community filtrates as the only carbon source, to its OD_620_ when grown on a synthetic glucose-M9 medium. **B**. Mean community function over generations. Green circles represent the mean functions of 92 cross-feeder communities (± 1 SD) in the selection line. Red circles represent the random control line. Asterisks indicate generations in which the community functions differ significantly between selection and control lines (Welch’s two-sample t test. *** indicates p < 0.001, ** indicates p<0.01, * indicates p<0.05). **C**. Maximal community function. Green triangles represent the mean of the triplicate measurement of the best performed community (± 1 SE) in the selection line. Red triangles represent the control line. Asterisks denote the same significant levels as those shown. **D**. Distribution of community functions in the selection and control lines over the seven generations. Bins are semi-transparent to show the overlapped counts in each line. **E**. Community functions of parent-offspring pairs in the selection line. Light red triangles denote the lineage of the dominant community that eventually spread over, dark red squares denote the lineage of the second dominant community (the last community being dropped), and the black circles represent the other lineages. Error bars are the standard errors of mean in the triplicate measurements. Grey dashed line represents the mean fitted linear regression from 1000 bootstraps. P-value reported here represents the fraction of non-significant linear regression slopes in 1000 bootstraps, e.g. p<0.001 means that linear regression of every bootstrapped data gives a significant slope (see Methods). The regression slope is an estimate of the trait heritability at the community-level (Goodnight 2000; Blouin et al. 2015). **F**. Muller plot showing the fraction of the metapopulation that is made up by the “descendants”of the different starting communities. The plot shows the rapid “fixation” of the dominant, highest function community. Each color represents one of the 12 soil and leaf samples from which the cross-feeder communities were initially inoculated. **G**. Heritability (h^2^) and mean function (± 1 SE) of the selection line at different rounds of selection (generations). The mean function is the as shown in **A**. The heritability drop after the fourth selection round coincides, as expected, with the lack of further response of the mean to selection.

Following previous work (Goldford et al. 2018; Lu et al. 2018) we started the experiment by suspending 12 different leaf and soil samples in M9 minimal medium, thus creating a heterogeneous set of starting inocula. Each inocula seeded seven or eight unique habitats (92 wells of a deep-well 96-well plate, each containing 400uL of M9 citrate media. The four additional wells are left as controls), as described in Methods. The plate was incubated at 30°C without shaking for 48hr. At the end of that period, a sample from each community was extracted and filter-sterilized, and the leftover cultures were stored in the refrigerator at 4°C. A volume of 50uL of the filtrate of each community, supplemented with 50 uL of 2X M9 salts, was added to a different replicate *E. coli* culture as the only carbon source. The cultures grew on a 384-well plate for 24hr at 30°C without shaking. The yield of the culture was determined by spectrophotometry, and normalized by the yield of an *E. coli* culture in the same plate and using M9 glucose as the only carbon source. The 23 communities whose filtrate had yielded the highest *E. coli* growth were selected, and used as the inocula to seed 92 new habitats in exactly the same manner as we did before. This artificial selection process was repeated for seven generations. Similar to previous studies (Swenson et al. 2000a,b) we generated a random selection control line, where each generation we selected 23 random communities to inoculate a new set of habitats. Other than that, the control experiments were performed in the exact same manner and using the same media and incubator as the artificial selection experiments (Methods).

We first quantified the mean function in the meta-community in each round of selection (generation), which we plot in Fig. 2B. The mean function of the random-selection control remained low throughout the experiment. By contrast, the mean function of the artificial selection line exhibited a strong initial response to selection, increasing by five-fold over the first three rounds of selection and leveling off after the fourth generation. At the same time, and despite this strong response of the mean, we also found that the function of the best performing community in the population did not improve in response to selection, remaining high but approximately constant over the experiment (Mann-Kandell test *τ* = −0.24, P = 0.56) (Fig. 2C).

When we plotted the distribution of community-functions at different generations, we found that the initial distribution (i.e. for the communities that were seeded from the heterogeneous set of soil inocula) was long tailed (Fig. 2D). This heritability of this community-level trait is initially very high (h^2^>0.85 over the first two rounds of selection, p<0.001) (Goodnight 2000): offspring communities that were seeded from a high function parent also had a high-function in the next generation, whereas those seeded from a low function community remained low in the next generation (Fig. 2E). As community-level selection proceeded, the metacommunity became more functionally homogeneous as the fraction of descendants from the two high-function initial inocula grew to take over the population (Fig. 2E). By the fifth generation, all of the communities were “descendants” of the two high-function initial inocula, and in the sixth generation a single soil inocula dominated (Fig. 2F). Once the population is entirely made up of replicates of the top initial community and the initial functional variance is entirely “spent” (Fig. 2G), the response to selection stops: the heritability h^2^ plummets, and becomes non-significant in the last three selection rounds (Methods).

## Discussion

We have presented the results of two artificial selection experiments on bacterial communities that utilized the “propagule”strategy to attempt to select for an optimized community function. The two experiments have important differences. However, they both highlight some of the main limitations of propagule based selection, and both illustrate the critical role of (heritable) community-level variability in artificial community-level selection (Goodnight 2000). In particular, we expect from the breeder’s equation that when the amount of heritable phenotypic variation in the population is high, the response of the mean trait to selection will be strong, whereas when it is negligible, no response will be observed.

One critical difference between the two experiments is the degree of functional variability in the starting community. When we designed the first experiment, part of what we wanted to learn is to what extent functional variability in the metacommunity could be supplied by endogenous processes such as spontaneous mutations or random transitions in ecological states. To that end, we seeded all 24 initial communities from the same synthetic, low diversity initial pool which consisted of four strains. We found that the heritability was not significant, and consistent with that finding neither the mean nor the maximum community function in the selection line improved in response to selection. Yet, both were higher in the artificial selection line than in the random selection control at the end of the experiment. This relative difference between the experimental and control lines was caused by a decline in the function of the random selection control during the stabilization period. This outcome is in line with previously reported results (e.g. (Swenson et al. 2000b)), where the effect of artificial community-level selection was more apparent in relative terms (i.e. by comparison to a random selection control) than in absolute terms (i.e. as marked by an increase in function relative to the ancestral community). Our interpretation is that the main effect of artificial selection in these experiments was to purge communities with low function (Rainey and Quistad 2020), rather than to improve those communities for which the function was already high.

We found this information helpful when we designed the second experiment. To stimulate a strong response to selection, we started the metacommunity with high initial functional and compositional variance, by seeding the communities from twelve different soil inocula. This design worked, and we found that the heritability was initially high (h^2^>0.85, p<0.001 over the first three generations). The large initial variance in community function fueled the strong response to selection observed in the first few rounds of selection. The increase in mean function ended when this variance was spent (the heritability h^2^ was not statistically significant after the third selection round, P>0.05), and all of the communities in the population were descendants of the highest performing community in the initial pool (Fig. 2F-G). The maximum function in the community did not increase and remained approximately constant over time. This finding indicates that the main effect of artificial selection was again to purge the low-performance communities, rather than improving the function of the highest performers. This suggests that the artificial selection protocol we implemented would have not worked better than simply taking the best performing community at the beginning of the experiment, and propagating it without selection (i.e. a simple exercise of bioprospecting at the community-level, or “eco-prospecting”)

For both of these experiments, it is reasonable to speculate that an improvement in the community-level function is likely to align against selection at the individual (cellular) level. For instance, a mutant with a higher amylase production than the ancestral population may engage in an ecological public-goods evolutionary game (Hauert et al. 2006, 2008; Rauch et al. 2017), and be selected against due to the higher cost of amylase production (West et al. 2006; Harrington and Sanchez 2014). Similarly, in the second experiment, mutants that appear in a population which are capable of using the metabolic byproducts secreted by other species in the community may in principle have a growth advantage, by being able to expand into open niches. This would in turn deplete those resources, lowering the amount of cross-feeding towards *E. coli*. We would thus anticipate that evolution is likely to erode both of those functions over time. Such potential conflicts between individual level selection and desired community-level functions are recognized as important hurdles for the engineering of microbial consortia for biotechnological purposes (Goldman and Brown 2009; Escalante et al. 2015; Xie et al. 2019).

Our results hint at some of the limitations and strengths of the propagule selection strategy. On the strengths side, we find that community traits are reliably passed from parent to offspring communities when we propagate the best performing communities without mixing or adding new migrants. Additionally, we find in both experiments that the propagule strategies are efficient at purging from the population those communities whose function has degraded. On the weaknesses side, the propagule strategies failed to improve the function of the best performing communities in our experiments. We hypothesize that this may be due to the lack of new variation entering the population through mutational processes alone, which is thus unable to overpower the non-heritable variation. The latter may arise from multiple sources, including measurement error when we quantify the function of each community, sampling stochasticity when we inoculate the offspring communities, and experimental day-to-day variation in media preparation, temperature fluctuations in the incubator, and other experimental artifacts (Xie et al. 2019)

Our experiments have several limitations, which may limit the scope of our findings. These shortcomings are shared with many other published experiments (Swenson et al. 2000b; Panke-Buisse et al. 2015). First, we only had one artificial selection and one random selection line in each experiment. These two experiments are very labor intensive and it would have been logistically challenging to even double the number of lines in both cases. Yet, it has been rightfully argued elsewhere that a lack of replication is a major problem of artificial selection experiments (Mueller and Sachs 2015), and this critique would also extend to both experiments reported here. Investigating how the community composition varied between the selected and non-selected lines may have clarified the role of ecological versus evolutionary processes. We did not save those samples, so unfortunately this is beyond the scope of what is now possible. Despite these shortcomings, we believe we have drawn valuable lessons from both of these experiments and thus that they are worth reporting. In particular, the outcomes of both experiments are in reasonable agreement with, and can be explained from, qualitative expectations of standard breeding theory. Looking to the future, and given the considerable challenges of doing these experiments in high-throughput (but see (Blouin et al. 2015)), we suggest that simulations such as those performed elsewhere (Williams and Lenton 2007a,b; Doulcier et al. 2019; Xie et al. 2019) will be very useful to explore different selection regimes, and thus to extract generic conclusions. Such efforts will be critical in order to develop a Theory of artificial selection of microbial communities that can guide the design of new protocols and experiments.

## Methods

### Strains, reagents, and growth conditions for amylolytic communities

Bacterial strains were obtained from ATCC (Manassas, VA, USA): *B*. *subtilis* (ATCC 23857), *B*. *megaterium* (ATCC 14581), *B*. *polymyxa* (ATCC 842), *B. licheniformis* (ATCC 14850). Glycerol stocks for each strain were struck out on nutrient broth agar (3g beef extract; 5g peptone; 15g agar per 1L DI water) supplemented with +0.2% starch (w/v, soluble, Sigma-Aldrich). Individual colonies were used to inoculate starter cultures (1mL; overnight; 30C). OD620nm measurements (Multiskan Spectrophotometer; Fisher Scientific) of 100 uL of overnight cultures were used to prepare an initial inoculum pool by mixing all four strains at an equal ratio (as measured by OD620nm; target OD620nm = 0.025 for each strain/= 0.1 total OD) in 1xbSAM media (prepared as described in (Sanchez-Gorostiaga et al. 2019)).

To begin the amylolytic community selection experiment, all wells of a 24-well plate were filled with 900uL of newly prepared 1xbSAM and each well was inoculated with 100uL of the initial inoculum pool prepared above (initial OD620nm = 0.01). The plate was covered with aeroseal rayon film (Sigma) and grown for 48 hours at 30C without shaking. After that time, a 500uL volume of each culture was filtered-sterilized (0.2 um centrifugal filter; nylon membrane; VWR 82031-358) and assayed in technical triplicate for enzymatic starch hydrolysis (see measuring amylolytic function below). OD620 nm (100 uL; Multiskan Spectrophotometer; Fisher Scientific) was measured for the four communities with the highest amylolytic function and for four random; non-selected communities (chosen by random number generation in Excel). Each community was then used to inoculate 6 “offspring” communities in three, 24 well plates (one plate for selected propagules; one plate for non-selected propagules; one plate for non-inoculated media control; 1mL 1xbSAM per well; initial OD620nm=0.01 as calculated as dilution of measured OD620nm of previous day’s culture). Incubation, amylolytic assay, and inoculation of selected/non-selected offspring were repeated every 48 hours. Reagents used throughout the experiment were identical to those used in (Sanchez-Gorostiaga et al. 2019), and growth and assay media were prepared in the same manner as described therein.

### Measuring amylolytic function

A 500uL volume of culture from each well was filtered-sterilized and assayed in technical triplicate for enzymatic starch hydrolysis (see measuring amylolytic function below). Three technical assays for each supernatant were carried out in a 96 well plate (Corning 3596). Per reaction, 20uL of enzyme supernatant was added to 180uL of 0.1% w/v starch solution and incubated for T=10min at room temperature. The amount of starch degraded over that incubation time was quantified using the quantitative Lugol iodine staining method described in (Inaoka and Ochi 2011; Sanchez-Gorostiaga et al. 2019), and which was in turn adapted from the classic Fuwa method (Fuwa 1954). For each assay plate, a negative control for both M9 media and 1xbSAM was also measured in triplicate as well as a standard curve (0.2%; 0.05%; 0.0125%; 0.003125% starch in M9). The amylolytic function of each community was determined as the fraction of starch degraded in the above assay.

### Sample collection for cross-feeder community

12 soil or leaf samples were collected at 12 different sites in and around Yale University West Campus in West Haven, CT, USA, and Yale University Campus in New Haven, CT, USA. For each sample at one collection site, five leaves or five scoops of soil within one meter radius were collected and pooled into one sample using sterilized tweezers or spatula. One gram of collected sample was placed in a 15 mL sterile tube and resuspended in 10 mL of 1X phosphate buffered saline supplemented with 200 ug/mL of cycloheximide to inhibit eukaryotic growth. Samples were homogenized by vortexing and were allowed to sit for 48 hours at room temperature. The supernatant solutions were used as inoculum for later experiments (see section below).

### Habitat inoculation, media, and cross-feeder community growth

Communities were grown in 0.2% citrate-M9 medium, prepared as described in (Lu et al. 2018) from the same stocks and reagents. 500 uL of culture media was added to each well of a 96-well deep-well plate (VWR). The first cell passage was supplemented with cycloheximide at a concentration of 200 ug/mL. Starting inocula were obtained directly from inoculating 4 uL of the supernatant of initial microbiota solution to 500 uL of culture media. Each of the 12 environmental samples was dispensed into seven or eight wells on the initial 96-well plate on which four wells were left for blank as media control. Plates were covered with Aerogel film (VWR) and incubated at 30 °C for 48 hours without shaking, then each community was homogenized by pipetting up and down 10 times. This initial plate was then replicated into two plates by inoculating 4 uL of grown media into 500 uL of fresh media. The two replicate plates corresponded to the selection line and the control line in the first community generation of the selection experiment. Both plates underwent six rounds of selection under the same incubation conditions described above, but differed in the communities selected to be propagated in each generation. Community phenotyping and the selection scheme are described in the next section.

### Community phenotyping and selection scheme

In the first community generation, communities in the 96-well plates were grown for 48 hours in 0.2% citrate-M9 media. To obtain the supernatant for community phenotyping, 200 uL of the grown media was spinned down at 3000 rpm for 15 minutes in an Eppendorf 5810 tabletop centrifuge and its supernatant was filter-sterilized using a 0.2 uM membrane 96-well filter (VWR 97052-096). The rest of the unfiltered grown media was kept at 4 °C for up to 30 hours as the inoculum for the next generation. The filtrate of each community was mixed with 200 uL of 2X M9 salts to make 400 uL of 1X M9 medium with the filtrate as the only source of carbon. 100 uL of this filtrate-M9 medium was dispensed to three wells of a 384-well deep-well plate (VWR), which enables us to measure the community function in N=3 replicates. To 20 additional wells, we added 100 uL of 0.2% glucose-M9 medium as a positive control. The border wells of the 384-well plate were filled with water and not used for cell culture in order to reduce the edge effect caused by evaporation. To all the wells with media, we inoculated 2 uL of an overnight pre-culture of *E. coli* (strain nohQ::Kan deletion strain from the Keio collection (Baba et al. 2006), derived from JW1541), which had been pre-grown for 24 hour in 2 mL of glucose-M9 medium at 37 °C in a 50 mL tube with constant shaking. The 384-well plate was then incubated at 37 °C for 24 hours without shaking, and then the optical density (620 nm) of the grown cultures was measured using a plate spectrophotometer (Fisher Scientific Multiskan Spectrophotometer).

The function for each community was determined as the ratio of the mean OD_620_ of the community filtrates, to the mean OD_620_ for the glucose-M9 cultures. In that time, the 96-well plate with communities was taken out of the refrigerator. The top 23 communities used to inoculate the next passage were determined by using a custom R script (Methods), which computed the community functions from input OD_620_ reads, ranked the communities according to their functions, and returned a list of 23 selected communities. These 23 communities were the parental communities and each of which was used to inoculate four offspring communities in the next generation. The same procedures were implemented on the control line in which the only difference was that random 23 communities, instead of the top communities, were chosen to propagate communities of the offspring generation. At generation four, an unexpected cross-contamination occurred in one of the four citrate-M9 blank wells on the selection line plate and it was accidently selected to propagate the later generations due to its high performance. Because we cannot trace the ancestral community of this lineage, we removed it from the analysis.

### Bootstrapping in heritability estimation

In order to take into account that the community function was measured in three technical replicates, we performed a bootstrapped linear regression to estimate the heritability for the selection line in the second experiment. To also track heritability as selection proceeded, heritability was computed from the functions of parent-offspring pairs within a selection round. For each parent-offspring pair, we randomly sampled the parent and offspring functions from their respective triplicate measurements. We then performed a simple linear regression, and recorded the significance level of its slope. This procedure was repeated for 1000 times (bootstraps) for each selection round. When more than 95% of the bootstrapped linear regression returns a statistically significant slope, we reported the mean of the bootstrapped slopes as the estimated heritability. Otherwise, we set h^2^=0.

## Data storage

All data in this paper and all data analysis scripts are publicly available and stored at: https://github.com/Chang-Yu-Chang/farmer_ecoli

## Author Contributions

CYC performed experiments, analyzed data, and wrote the paper. MLO performed experiments and analyzed data. DB designed and performed experiments. AS conceptualized the project, designed experiments, analyzed data, and wrote the paper.

## Acknowledgments

We want to thank Maria Rebolleda-Gomez, Sylvie Estrela, Jean Vila and all other members of the Sanchez lab for their input and comments on the manuscript. The funding for this work partly results from a Scialog Program sponsored jointly by Research Corporation for Science Advancement and the Gordon and Betty Moore Foundation through grants to Yale University (AS). This work was also supported by a young investigator award from the Human Frontier Science Program (RGY0077/2016) and by the National Institutes of Health through grant 1R35 GM133467-01 to AS. CYC was supported by a graduate fellowship from the Taiwanese government (Government Scholarship to Study Abroad).

## Notes

### Competing Interest Statement

The authors have declared no competing interest.

